# Distinct mesoscale cortical dynamics encode search strategies during spatial navigation

**DOI:** 10.1101/2023.03.27.534480

**Authors:** Daniel Surinach, Mathew L Rynes, Kapil Saxena, Eunsong Ko, A David Redish, Suhasa B Kodandaramaiah

## Abstract

Spatial navigation is a complex cognitive process that involves neural computations in distributed regions of the brain. Little is known about how cortical regions are coordinated when animals navigate novel spatial environments or how that coordination changes as environments become familiar. We recorded mesoscale calcium (Ca^2+^) dynamics across large swathes of the dorsal cortex in mice solving the Barnes maze, a 2D spatial navigation task where mice used random, serial, and spatial search strategies to navigate to the goal. Cortical dynamics exhibited patterns of repeated calcium activity with rapid and abrupt shifts between cortical activation patterns at sub-second time scales. We used a clustering algorithm to decompose the spatial patterns of cortical calcium activity in a low dimensional state space, identifying 7 states, each corresponding to a distinct spatial pattern of cortical activation, sufficient to describe the cortical dynamics across all the mice. When mice used serial or spatial search strategies to navigate to the goal, the frontal regions of the cortex were reliably activated for prolonged durations of time (> 1s) shortly after trial initiation. These frontal cortex activation events coincided with mice approaching the edge of the maze from the center and were preceded by temporal sequences of cortical activation patterns that were distinct for serial and spatial search strategies. In serial search trials, frontal cortex activation events were preceded by activation of the posterior regions of the cortex followed by lateral activation of one hemisphere. In spatial search trials, frontal cortical events were preceded by activation of posterior regions of the cortex followed by broad activation of the lateral regions of the cortex. Our results delineated cortical components that differentiate goal- and non-goal oriented spatial navigation strategies.

## INTRODUCTION

Successful navigation to a goal can be achieved using multiple behavioral strategies. Evidence suggests that mammals (including mice, rats, cats, dogs, monkeys, and humans) have access to multiple decision processes which are used at different times, and which can be separated out with appropriately defined behaviors^1–3^. The Barnes maze is a goal-finding task that rats and mice learn readily^4,5^. In this task, mice are placed at the center of a well-lit environment and try to find an escape goal. Initially, mice will have to search randomly as they learn the basics of the task itself, but with experience, mice learn to go directly to the goal. Mice who know the task objective (find the escape hole) but who do not know the spatial location of the goal perform a serial search through many potential hiding holes^6^.

The incoming sensory information during spatial navigation is processed widely across the cortex, and information about space has been shown to be present in many cortical regions ^7–9^. A number of cortical regions have neurons that reflect allocentric information ^10–12^, while other regions have neurons that encode turns and other egocentric information^11,13–16^, and specifically retrosplenial cortex has been shown to encode both egocentric and allocentric information ^17^. Neural activity in association areas of the cortex are also implicated in encoding landmarks ^18–20^, route planning ^11,13,21^ and associating allocentric cues with motor decisions ^22^. It has also been shown that interactions between brain regions are important for various cognitive and spatial navigation tasks ^18,23–25^.

While the different navigation strategies have been shown to reflect different computational processes in individual cortical and subcortical regions ^1,26–31^, it remains unknown how cortical signals are coordinated between different brain regions when these different strategies are being used for navigation. The measurably different strategies that mice show in the Barnes maze provide an opportunity to determine these cortical interaction changes that drive strategy changes.

We imaged calcium (Ca^2+^) activity across most of the dorsal cortex of freely behaving mice while they solved the Barnes maze. We found that consistent with previous literature, mice rapidly and progressively used less time to reach the goal with increasing experience. Using a novel clustering algorithm, we decomposed the cortical dynamics into a low dimensional common state space, with each state corresponding to a pattern of cortical activation. We analyzed the temporal sequences of state activation and found distinct sequences of state transitions in the early part of the trial when mice made decisions about the direction to approach the edge of the maze. These sequences of cortical state activations indicate distinct sets of brain wide circuits are engaged when different behavioral strategies are used to solve the maze.

## RESULTS

### Mesoscale calcium imaging in mice learning a 2D spatial navigation task

We imaged calcium activity across 8×10mm^2^ of the dorsal cortex, encompasses parts of the primary and secondary motor cortices, the somatosensory and barrel cortices, the retrospenial cortex and part of the visual cortex in both hemispheres using a miniaturized head mounted camera (mini-mScope, (Rynes & Surinach et al. 2021)) in eight freely behaving Thy1-GCaMP6f mice ^33^, as they solved the Barnes maze (**Fig. 1a-c**). As mice learned the location of the goal, they exhibited expected results in the strategies used to search for the goal, which could be categorized as random, serial, or spatial search strategies ^5^ (**Fig. 1d.** As trials progressed, mice demonstrated a reduction in primary errors, or the number of incorrect holes checked prior to reaching the correct location of goal, and primary latency, or the initial trial time until the goal location is found (**Fig. 1 e-f**).

**Figure 1:**
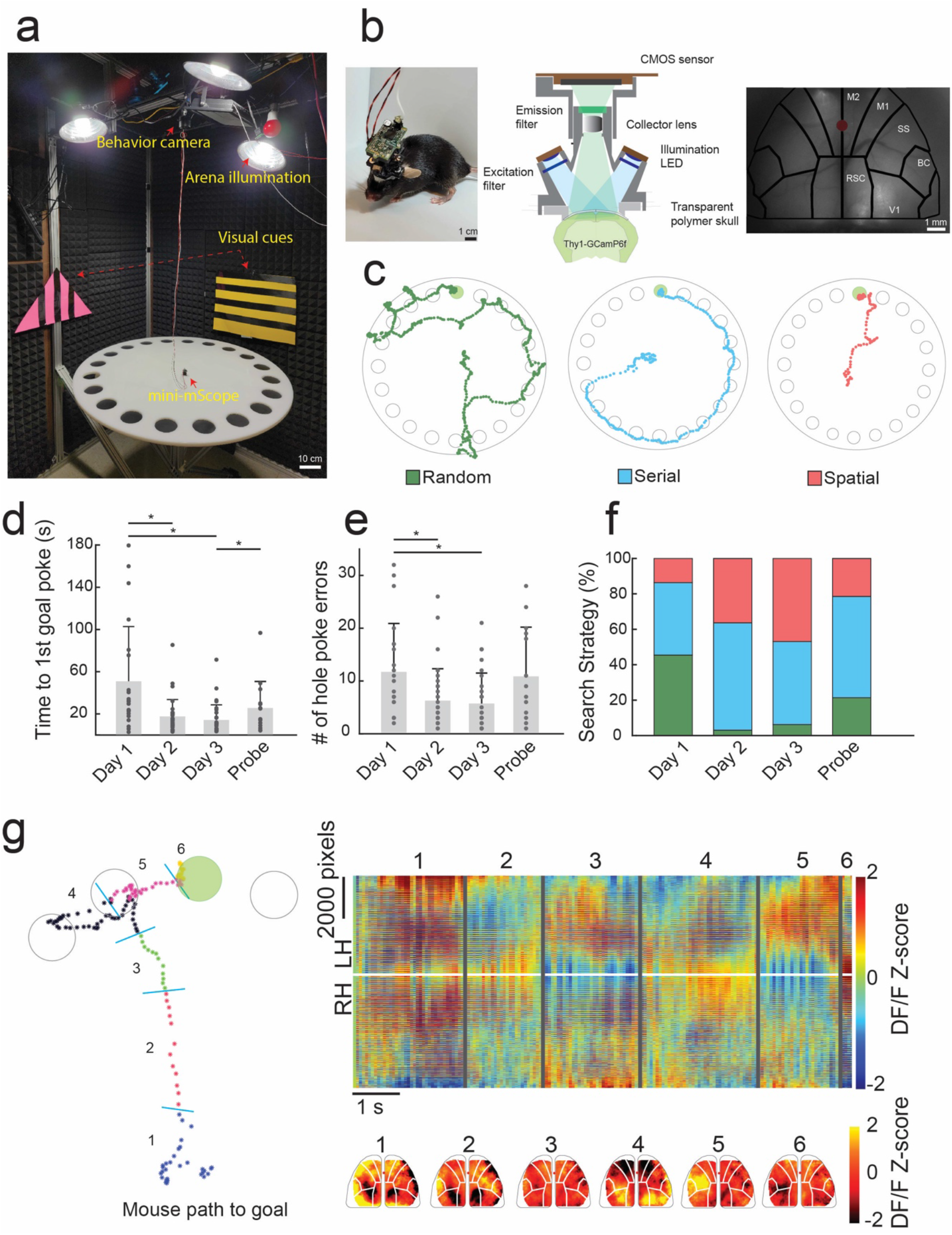
Mesoscale calcium imaging during spatial navigation. **a)** Photograph of the behavioral setup including the mini-mScope and the Barnes Maze. **b)** Left: Photo of a mouse bearing the mini-mScope in the behavioral arena in a). Middle: Computer aided design (CAD) cross-sectional view of the mini-mScope. Right: Photo of raw imaging field of view (FOV) through the mini-mScope with an Allen atlas overlaid. Red dot indicates bregma. M1- Primary motor cortex, M2 – secondary motor cortex, SS – somatosensory cortex, BC – barrel cortex, V1 – visual cortex, RSC – retrosplenial cortex. **c)** Traces obtained from tracking data of one mouse which utilized random, serial, and spatial search methods as it learned to navigate the Barnes maze. **d)** Bar plot showing the mean primary latency, or time to first goal hole discovery, across days as mice learned to navigate the Barnes maze. * indicates p < 0.05, Wilcoxon rank sum test. **e)** A bar plot showing the mean number of primary errors, or the number of times the mouse checked an incorrect hole before reaching the goal. * indicates p < 0.05. **f)** A bar plot showing the percentage of search strategies utilized across all trial days. **g)** *Left:* Tracking data from a spatial trial in which the mouse makes a single error on the way to the goal. The trace is annotated with periods that correspond to state-like shifts in calcium data across the cortex shown in the graph on the right. *Right:* a map of the calcium data across the entire FOV acquired during the trial shown in the left panel. Numbered lines correspond to state-like global calcium activity transitions observed during the behavioral periods marked in the left panel. Pseudo color maps of the calcium DF/F z-score from frames during each behavioral period are shown below. All error bars indicate sample standard deviation.

Primary latency decreased from 51.0 ± 51.7 s (55.1 Interquartile range, IQR) on day 1 acquisition trials to 17.6 ± 16.1 s (11.0 IQR) on day 2, and 14.2 ± 14.4 s (11.1 IQR) on day 3 (**Fig 1d,** Day 1 vs Day 2 p = 0.0006, Day 1 vs Day 3 p = 0.0001, Day 3 vs Probe p = 0.044, Wilcoxon ranked sum test). The primary latency increased to 26.0 ± 24.7 s (21.03 IQR) s when the goal location was moved on the probe trial, where the location of the goal was altered. Similarly, the number of primary errors decreased from 11.7 ± 9.2(11 IQR) on day 1, to 6.3 ± 6.1(7 IQR) on day 2 and 5.7 ± 5.7(7.5 IQR) on day 3 across all mice, and the number of primary errors increased to 10.9 ± 9.2(16 IQR) when the goal location was changed in the probe trial (**Fig. 1e,** Day 1 vs Day 2 p = 0.012, Day 1 vs Day 3 p = 0,0001, Wilcoxon ranked sum test). These results are consistent with previous results obtained in this task^21^, indicating mounting the mini-mScope did not interfere with behavior.

Across trials, mice utilized increasingly non-random search methods as they learned to navigate the maze. On day 1, 54.5% of trials were nonrandom, whereas 45.5% were random. On day 3, 93.7% of trials were non-random and 6.3% of trials were random. As trials progressed 13.6% of trials were spatial on day 1, 36.36% of trials were spatial on day 2, and 46.8% of trials were spatial on day 3 (**Fig 1f**).

While the white noise and bright lights were presented as mildly noxious stimuli and motivated the mice to navigate to the goal progressively faster, mice rarely entered the goal immediately after first poke (21% of trials, n=13/63 trials), preferring to explore the arena. In a subset of trials, mice explored the two nearest holes and the edge around the goal hole in 32% of trials (n=20/63), entering the goal hole 5-30 s after nearby exploration. A large subset of mice (46%, n=29/63) chose to repeat one or more searches around the maze after first goal poke before entering at some later trial time. Thus, while the animals were motivated to go to the goal, the environment was not excessively stress-inducing such that mice were not prevented from exploring the maze further.

### Mesoscale cortical dynamics exhibited discrete shifts in cortical activation patterns

The mini-mScope imaged a field of view (FOV) of 8 mm x 11 mm, with a craniotomy encompassing 6 brain regions: primary motor cortex (M1), somatosensory cortex (SSC), premotor/frontal cortex (M2), retrosplenial cortex (RSC), primary visual cortex (V1), and barrel cortex (BC) on each hemisphere at a resolution of ~39-56 μm per pixel from the center to lateral edges of the FOV. As the mice navigated the maze, prolonged patterns of calcium activation across the FOV occurred sporadically, with shifts between these calcium activity patterns occurring at ~0.2-1 s time scales (**Fig. 1g**).

We used an image correlation and clustering methodology to cluster spatial pattens of calcium activity observed in individual frames into groups of highly correlated images with similar patterns of cortical activation. We refer to these groups of highly correlated images as cortical activation ‘states’ (**Fig 2a-b, Supplementary Fig 1a**). Briefly, the z-scored calcium DF/F activity recorded at each time frame was correlated with every frame recorded for a mouse across all trials, forming an image correlation matrix. The data in this matrix was then iteratively clustered into increasing numbers of states. The number of states needed to optimally cluster the cortical activity patterns is not known *a priori.* We used a t-distance optimization algorithm to determine the optimal number of states that could segregate the image correlation matrix into groups to maximize the correlations between images within a group while simultaneously minimizing the correlations between images across groups ^34^ (See Methods and **Supplementary Fig. 1** for more details). We found that 5-10 states optimally described calcium activity clusters across each mouse (**Supplementary Fig 1b, Supplementary Fig 2**). An example of this clustering methodology for one mouse is shown in **Figure 2a-b**.

**Figure 2:**
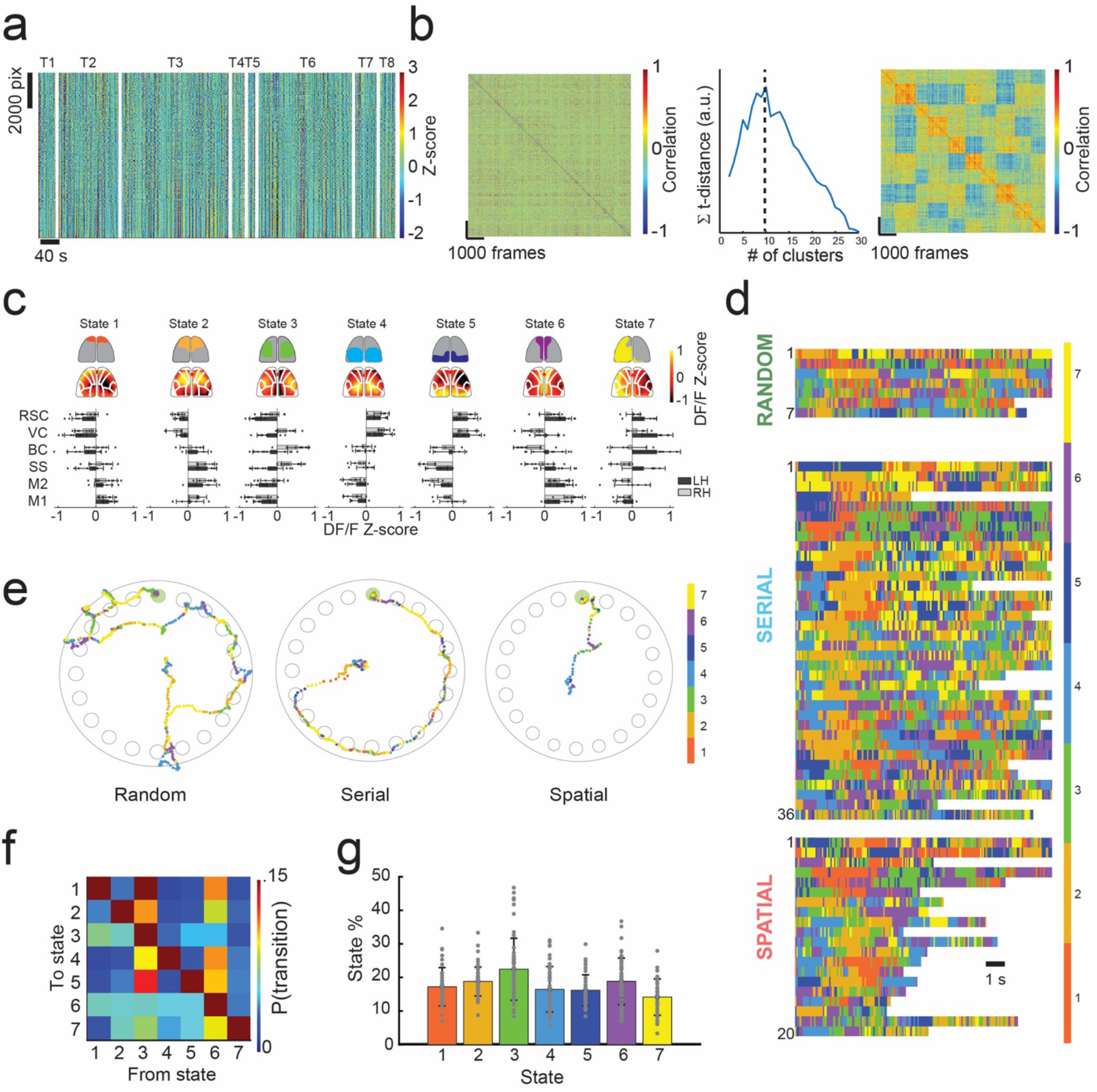
Identifying brain states from mesoscale calcium activity. **a)** Example of the method used to identify cortex-wide brain states from widefield calcium imaging during spatial navigation from one mouse. Data from all trials for one mouse is shown. All pixels across the FOV are plotted vs time. Trials are indicated by T1-T8 labels, separated by the white lines. **b)** *Left:* A correlation matrix is constructed by computing the image correlation between all frames. K-means clustering is used to organize the correlation matrix into highly correlated groups, denoted as states. *Center:* The number of states is determined by using an optimization algorithm which maximizes intra-cluster correlation while minimizing inter-cluster correlations. The maximum t-distance value indicates the optimal number of states for this mouse (k=10 states here). *Right:* the result of re-sorting the correlation map on the left into an optimized number of clusters determined with k-means clustering and t-distance optimization, resulting in 10 states for this mouse. **c)** Common state space model across n = 8 mice and 63 trials. Optimum number of states varied from 5-10 states across all mice, with an average of 6.4 ± 2 states (**Supplementary Figs. 1** and **2**). 7 states were selected as sufficient to describe the state space across mice. States were identified by cortical areas across the FOV with high DF/F z-score calcium signal. The top row illustrates simplified activity maps with high DF/F z-score activity. Below the top row are average DF/F z-score heat maps for the mouse in **a-b** which fit into the common state space. Bottom: bar graphs depicting the average DF/F z-score of cortical regions across mice using the Allen atlas. **d)** Time series of state activation of the first 12 seconds of all trials plotted in a time series. Color bar indicates state number. The top row are random trials, the middle row are serial trials, and the bottom row are spatial trials. **e)** Examples path plots of random, serial, and spatial trials with state number overlayed on mouse tracking data. Color bar indicates state number. **f)** State transition probability matrix across all trials. **g)** Bar graph of the total state activation probabilities across all trials.

To identify a common state space to describe activity in all mice, similar clustering methodology was employed. Briefly, the average DF/F activity for each state identified per mouse was calculated by averaging activity across all frames within each state. The average frames for each state for all mice were then correlated to form a second image correlation matrix across all mice (51 x 51 matrix, **Supp. Fig. 1a**). The image correlation matrix was then sorted into 7 states via k-means clustering to construct the intra-mouse state space model.

The spatial distribution of the mean calcium activity of all seven states for one mouse is shown in **Figure 2c top**. Additionally, a bar graph of the mean DF/F activity patterns for each ROI in the Allen brain atlas across all 7 states in each mouse (**Fig 2c, bottom)**. States 1 and 2 were characterized by high calcium activity in the frontal regions of the FOV. State 3 was characterized by high activity in several cortical areas of each hemisphere, with peak activation in bilateral somatosensory, primary motors, and antero-lateral retrospenial cortex. States 4 and 5 were characterized by high calcium activity in the posterior regions of the FOV. State 6 was characterized by high calcium activity in the vicinity of the midline. Lastly, state 7 was marked by activity distributed broadly across the left hemisphere. Observed mean activation patterns for states 1-6 were lateralized in most mice (**Supplementary Fig. 2**), perhaps indicating functional specialization between the cortical hemispheres during navigation.

Every mouse had one of state 1 or 2 present where frontal regions of the cortex were active, with n = 4 mice expressing both states. Additionally, every mouse had one of state 4 or 5 present, where the posterior regions of the cortex were active, with n = 2 mice exhibiting both states. State 3 and 6 where the lateral regions of the cortex and the medial regions of the cortex were respectively active were present in all mice (n = 8), and state 7, where the activity was higher in predominantly in the left hemisphere was present in n = 5 mice (**Supplementary Fig. 2**). Example montages of DF/F z-score activity for commonly occurring state transitions are shown in **Supplementary Figures 3** and **4**. The time series of detected states during the first 15 seconds of each trial is shown in **Figure 2d**, where rows denote trials for each search strategy, and colors signify the state that each frame in that trial was assigned. White spaces denote the trial has ended when the mouse enters the goal hole. Examples of the state activation along the path taken by a mouse during a random, serial, and spatial search trials are shown in **Figure 2e**. Similar visualization of state activation along the paths traversed by the mice in all trials are shown in **Supplementary Figures 5-8**.

We evaluated the probability of a particular cortical activation state being active. For all states, mean state activation probability varied between 14.2% - 22.7% (**Fig. 2f-g**). States 3 and 6 which are present in all the mice had slightly higher activation probabilities of 22.5 ± 9.2 % and 18.8 ± 6.9% respectively. Thus, there was no one state having a dominant activation probability. Grouping trials by search strategy (**Supplementary Fig. 9a**), we observed no significant differences in state activation probabilities for any of the states. The mean state activation probabilities did not change substantially as mice performed successive trials (**Supplementary Fig. 9a right**).

We further examined how cortical activation changed from one state to the other by constructing state transition probabilities matrices for serial search trials and spatial search trials (**Supplementary Fig. 9b**). Notably, state 3 had a high probability of 18.7% and 15.3% to transition to state 1 in random and serial trials, respectively. Transition probability from state 3 to state 1 in corresponding spatial trials decreased to 6.3% during spatial trials. State transition probabilities from state 5 were low (<6%) when transitioning to other states in trials on which mice used a random search strategy. In trials on which mice used a serial search strategy, state 5 transitioned to state 6 with a probability of 6.1%. In contrast, state 5 transitioned to state 3 and 7 with probabilities of 6.3% and 8.7% respectively during trials which mice used a spatial search method. These results highlight how cortical dynamics were different for the trials with different behavioral strategies.

### Frontal regions of the cortex are activated for prolonged durations shortly after trial initiation

Representing the patterns of cortical activation in a low-dimensional state space allowed us to examine trial-by-trial variation in cortical dynamics during the spatial navigation task. We observed repeated temporal sequences of state activation that occurred shortly after trial initiation. Trials typically started with a variegated sequence of states, but then transitioned to a clear and prolonged period of activation of the one or both frontal cortex active states (states 1 or 2) near the start of the trial (**Fig. 2d**). These prolonged durations of frontal cortex states (henceforth referred to as frontal state activation event or FSA event) could be algorithmically identified as conditions where state 1 or 2 was active for more than 1 second near the start of the trial (**Fig. 3a**). The FSA events occurred in 57.1% of trials where mice used random search method, 91.7% of trials which the mouse used a serial search method, and 85.0% of trials where the mouse used a spatial search method (**Fig. 3b**). These FSA periods were primarily associated with non-random search strategy trials. Overall, mean onset to the FSA event was 2.3 ± 1.9 s. In trials in which mice performed a random search strategy, the mean onset to the FSA event was 1.4 ± 1.2 s, whereas in serial search trials it was 2.3 ± 2.0 s, and 2.5 ± 2.0 s in spatial search strategy trials (**Fig. 3c,** p = 0.46 random vs serial, p = 0.30 random vs spatial, p = 0.45 serial vs spatial). The mean duration of the FSA event was 2.0 ± 0.7 s. The duration of the FSA event at the beginning of trials were also longer in serial search and spatial than in random strategy trials. In trials on which the mice performed a random strategy, the mean duration of the FSA event was 1.5 ± 0.4 s, whereas it was 2.0 ± 0.6 s in serial trials, and 2.2 ± 1.0 s in spatial trials (**Fig. 3d,** p = 0.082 random vs serial, p = 0.18 random vs spatial, p = 0.98 serial vs spatial).

**Figure 3:**
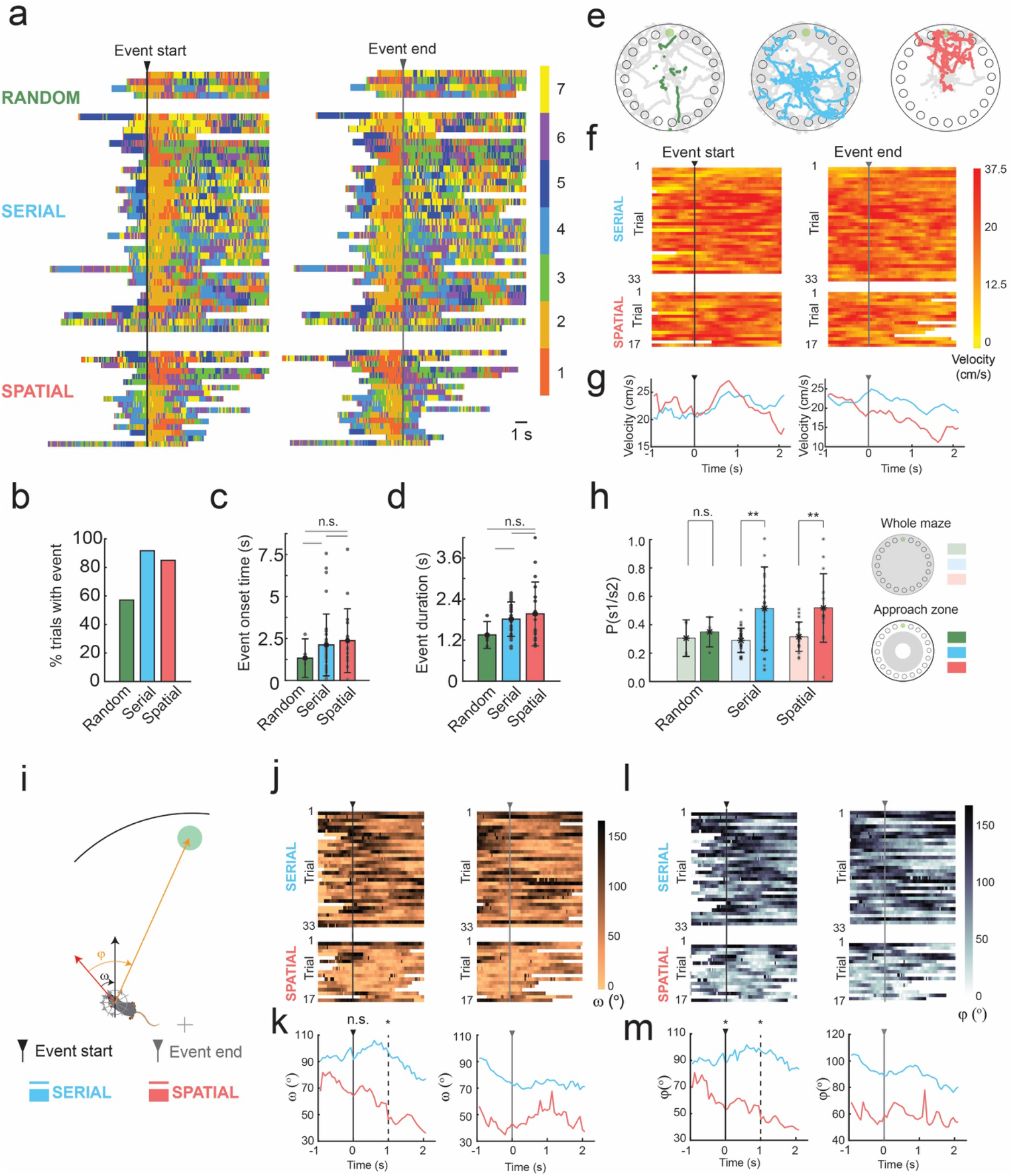
Prolonged activation of frontal states at trial start. **a)** Time series of state activation of all trials containing >1 second activation of frontal states 1 or 2 aligned to the start of the frontal state activation event (FSA event, left) and the end of the FSA event (right) **b)** A bar graph depicting the total percentage of trials in which FSA event occurred when mice used random, serial and spatial search strategies. **c)** A bar graph depicting the mean onset time to the FSA event for trials which the mouse utilized each search strategy. **d)** A bar graph depicting the total duration of the of the FSA event for trials which the mouse utilized each search strategy. **e)** Mouse tracking data from trials in which the mouse utilized random, serial, and spatial search strategies. Colored points indicate the frontal state activation period and gray points indicate the rest of the tracking data to first goal poke. **f)** Plots depicting the velocity of mice in trials with a FSA event. The FSA event period is aligned to the start and end of the event for serial and spatial trials. **g)** Average velocity plots made from **f** with serial and spatial trials superimposed before and after the frontal state activation event. **h)** State activation probability of frontal state 1 or 2 across the whole maze and within the approach zone. ** indicate p<0.001, Wilcoxon rank-sum test. **i)** Schematic of allocentric and egocentric head angles, ω and φ, respectively**. j)** Allocentric head direction plots of trials with the frontal state activation period aligned to the start and end of the event for serial and spatial trials. **k)** Average allocentric head direction plots made from **j** with serial and spatial trials superimposed before and after the frontal state activation event. * indicates p < 0.05 (p = 0.104 at 0s, p = 0.002 at 1s after FSA initiation, Wilcoxon rank sum test). **l)** Egocentric head direction plots of trials with the frontal state activation period aligned to the start and end of the event for serial and spatial trials. **m)** Average egocentric head direction plots made from **l** with serial and spatial trials superimposed before and after the frontal state activation event. * indicates p < 0.05 (p = 0.02 at 0s, p = 0.002 at 1s after FSA initiation, Wilcoxon rank sum test).

### Frontal state activation events coincided with approach to edge of the maze

We next evaluated the behavior of the mice around the FSA events in serial and spatial search trials by examining the position, velocity, and head direction of the mice (**Fig. 3e-l**). Plots of the location of the mice during the FSA event indicated that the FSA event occurred when mice approached the edge of the maze from the initial starting location at the center of the maze (**Fig. 3f**). In 84.4% of trials with a FSA event, the event initiated before or during the mouse’s approach to the edge of the arena. The FSA event began during the initial period of the trial before the mouse approached the edge and as it investigated its surroundings. The FSA events occurred before the mouse reached the edge of the maze in 60.6% of trials with an FSA event for serial trials vs 35.3% of trials for spatial search strategies with an FSA event. The FSA events were also accompanied by an increase in velocity of the animal, with instantaneous velocity peaking ~800ms after event onset in both serial (mean peak velocity of 25.5 cm/s, **Fig. 3f top left** and **Fig. 3g left**) and spatial trials (mean peak velocity of 27.1 cm/s, **Fig. 3f bottom left** and **Fig. 3g left**). The end of the FSA event coincided with a decrease in velocity in both serial and spatial trials, with a steeper decline in velocity in spatial trials starting 50 ms prior to the end of the event as mice approached the vicinity of the goal (**Fig. 3f right** and **Fig. 3g right**). Correspondingly, the probability of activation of either one of the frontal states was significantly higher in the approach zone of the maze, as compared to state activation probabilities across the whole maze in serial and spatial trials (p = 0.0022 serials trials, p = 0.003 spatial trials, Wilcoxon rank sum test).

### Frontal activation state events in spatial trials followed initial orientation towards the goal

The period before the FSA event is likely a self-localization event in which mice survey the space before deciding on direction of approach to the edge of the mice. We examined the changes in both the allocentric heading direction angle (ω), and the egocentric heading direction angle (φ) of the mice at the start of the FSA event (**Fig. 3i-m**). When mice employed spatial search strategies, mice oriented towards the goal quadrant (I ω I < 45°) in 59% of trials (10/17 trials) at the onset of the FSA event, with an increased fraction (76.4%, 13/17 trials) 500 ms after event onset. In contrast, mice were oriented towards the goal quadrant in the allocentric reference frame in only 27% of serial trials (9/33) at the event onset (**Fig. 3j top left**). Mean ω was 64 ± 62° in spatial trails as compared to 92 ± 50° in serial trials at FSA event onset (p = 0.103, Wilcoxon rank sum test). Mean ω was 46 ± 43° in spatial trails, significantly lower as compared to 97 ± 45° in serial trials 1s after FSA event onset (p = 0.017, Wilcoxon rank sum test).

Similar differences in egocentric heading direction angles between serial and spatial trials at the start of the FSA event (**Fig. 3l and m**). When mice employed spatial search strategies, mice oriented in the direction of goal quadrant (I φ I < 45°) in 65% of trials (11/17 trials), with an increased fraction (82%, 14/17 trials) 500 ms after event onset. In contrast, mice were oriented towards the goal quadrant in the egocentric reference frame in 24% of serial trials (8/33) at the event onset, with no decline in egocentric heading direction angle observed after event onset (**Fig. 3l top left**). Mean φ at event onset was 63 ± 59°, significantly lower than the mean φ of 98 ± 46° in serial trials (p = 0.020, Wilcoxon rank sum test). Significant differences in mean φ were maintained 1s after event onset (**Fig. 3m left,** p = 0.020, Wilcoxon Rank-sum test).

### Sequences of state transitions before activation of the frontal cortex were search strategy dependent

We next evaluated if there were differences in sequences of state activation during specific periods around the FSA events. Examining state activation probabilities in the duration of time prior to the FSA event period revealed differences between serial and spatial search methods (**Fig. 4a**). In trials where mice utilized serial searches, state 7 had an activation probability of 29.3 ± 29.2% prior to FSA event. In spatial trials, the activation probability reduced to 14.1 ± 5.4%. State 3 had an activation probability of 28.7 ± 26.9% prior to the FSA event in spatial search trials, significantly higher than the activation probability in serial search trials (12.4 ± 19.6%, p = 0.007, Wilcoxon ranked sum test, **Fig. 4a**). State 6 activation probability did not change notably between the two search strategies (16.5 ± 17.4% serial search trials, 11.9 ± 13.6% spatial search trials). Examining state transition probabilities in the period before the FSA event revealed differences in dynamics of cortical activity between spatial and serial trials **(Fig. 4b)**. Most prominently, state 3 had a high probability of transitioning to many states in spatial trials, but not in serial trials.

**Figure 4:**
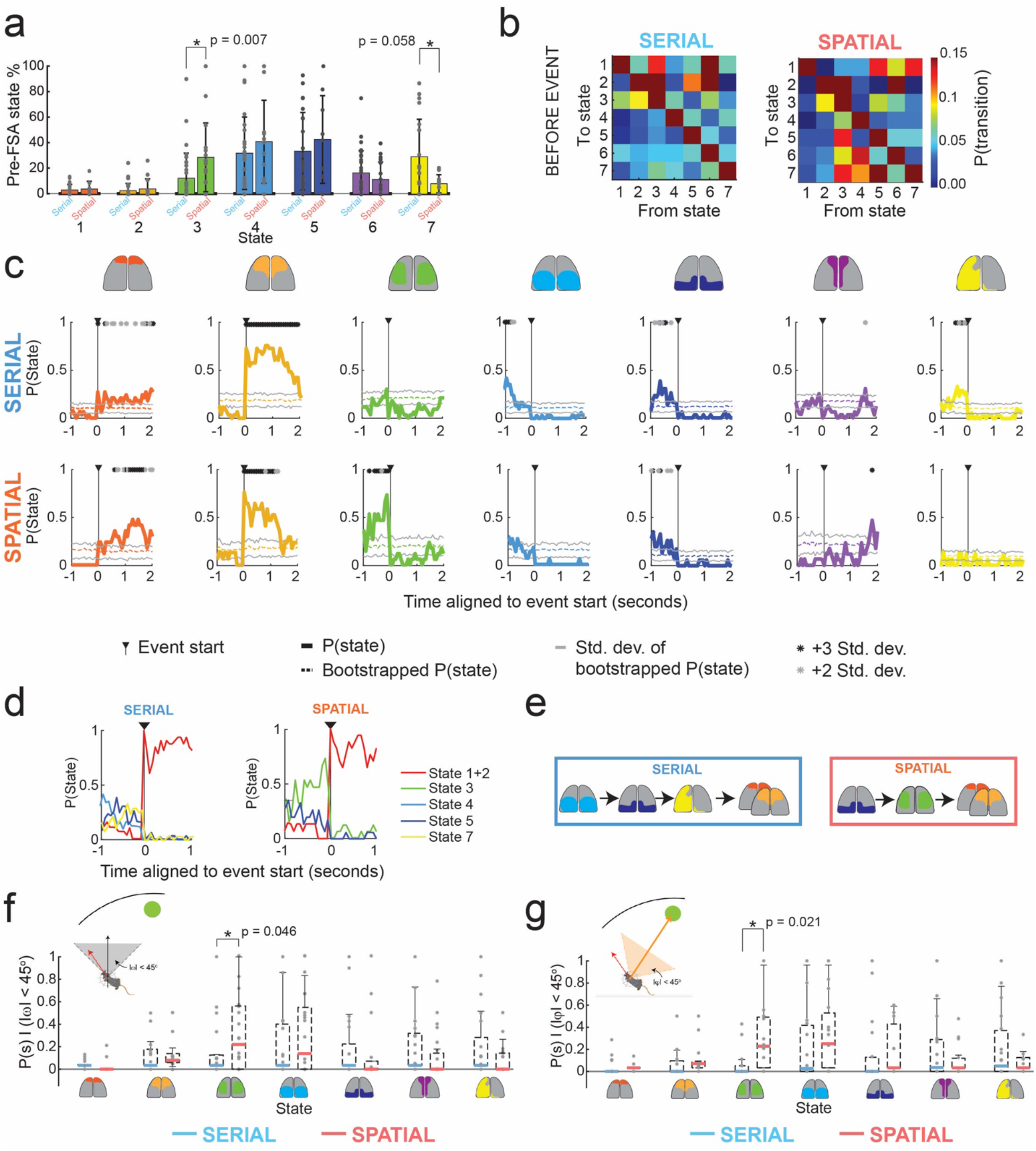
Sequences of states before FSA event depend on search method. **a**) Bar plots of states activation probabilities for all states before the FSA event in serial and spatial trials. Statistically significant differences are highlighted (Wilcoxon rank sum test) **b)** State transition probabilities before the FSA event for serial and spatial trials. **c)** Peri-event state probability histograms aligned to the start of the frontal state activation event in serial and spatial trials. Solid color lines denote the average peri-event probability across all trials. Transparent solid color lines indicate the average of 100 randomized bootstrapped trials with standard deviation lines in gray. Asterisks indicate statistical significance against bootstrapped data using an Anova test with a Bonferroni correction (gray p<0.05; black p<0.01). **d)** Summary of statistically significant probability of state activation in **g**. **e)** Simplified state transition schematic of cortical activation states 1 s before and during the FSA event using the statistically significant state activation periods found in **c** for serial and spatial trials. **f)** State activation probability box plots generated for allocentric head direction for |ω|<45 degrees for serial vs spatial trials. **f)** State activation probability plots generated for egocentric head direction for |φ|<45 degrees for serial vs spatial trials. Statistically significant comparisons are highlighted (Wilcoxon rank sum test).

To quantify the patterns of state transitions leading up to the FSA, we constructed peri-event state probability histograms (**Fig. 4g**). As a control, we generated randomized data by performing 100 bootstraps of the time series of states for each trial. We determined if a state’s activation probability was statistically significant from the bootstrapped trials by using an Anova test with a Bonferroni correction of the mean state activation probability aligned to the FSA period to the bootstrapped data. In serial trails, the 1 s period leading up to the FSA event was marked by significantly higher activation of state 4, followed state 5 and then by state 7 when compared bootstrapped mean. In contrast, in spatial trials, the same 1 s period was marked by activation of state 5 that was followed by activation of state 3 before entering the FSA period (**Fig. 4d**). These results indicate that the sequences of state transitions occurring before the FSA events were search strategy dependent (**Fig. 4e**).

### State 3 was preferentially active before FSA during goal-heading direction in spatial but not serial trials

When considering the entire duration before the FSA event, heading direction in the allocentric reference frame was significantly more aligned towards the goal in spatial search trials (77.1° ± 50.4°) as compared to serial search trials (94.9° ± 52.7°, p<001, Kruskal-Wallis test). Similarly, evaluating the egocentric heading direction revealed significantly more alignment towards the goal in spatial search trials (88.1° ± 50.3°) as compared to serial search trials (101.8° ± 51°, p<0.001, Kruskal-Wallis test). Thus, there was an overall change in tuning of heading direction for most states that differed between serial and spatial trials. We next asked if the animals head orientation affected the activation of states. We examined the times when head direction of the mice was aligned to the goal quadrant in the allocentric reference frame (I ω I < 45°, **Fig. 4f**) and egocentric reference frame (I φ < 45°, **Fig. 4g**). Within these events we asked what the likelihood of a certain state being active was prior to the FSA event. State 3, which was significantly more likely to be active immediately prior to FSA event onset in spatial trials (**Fig. 4a,c**), was much more likely to be active when animals were oriented towards the goal quadrant in the allocentric frame of reference during spatial trials), as compared to serial trials (mean P(s3) spatial = 0.33, mean P(s3) serial = 0.13, p = 0.046, Wilcoxon Rank-sum test). For egocentric goal orientation, state 3 also had a higher probability of being active while mice were oriented to the goal as well (mean P(s3) spatial = 0.31, mean P(s3) serial = 0.07, p = 0.021, Wilcoxon Rank-sum test). No significant differences were found for any of the other states. These results indicate that state 3 was preferentially activated when the animals head direction was oriented towards the goal in spatial trials, but not in serial trials. Thus, activation of state 3 in spatial trials may indicate a recognition of the goal direction in spatial trials when mice make direct approaches to the goal.

## DISCUSSION

We discovered coordinated sequences of brain-wide activity patterns reflected in mesoscale cortical activity on a spatial navigation task that differentiated goal-oriented and non-goal-oriented strategies. The clustering algorithm we developed in this study identified 7 cortical activation states that were generalizable across mice and trials, and 15 state transitions that occurred frequently during this spatial navigation task. Similar numbers of dynamic motifs have been independently described in studies looking at mesoscale calcium dynamics during head-fixed spontaneous behaviors ^35^, with distinct dynamics observed during memory guided and sensory guided tasks ^36^, and uninstructed movements during sensory decision making ^37^ and locomotion ^38^. These findings suggest that such generalizable repeated sequences of cortical activity may underlie a diverse set of behaviors. Our data show that these sequences differentiate decision strategies, most likely due to changes in the computations underlying these decision strategies.

Trial initiation was marked by an initial duration lasting 1-2 seconds of variegated sequences of states while animals were in the center of the maze near the starting location, followed by prolonged activation of states associated with activation of frontal areas of the cortex lasting 1-2 seconds (FSA event) as the animals turned towards the edge of the maze. Despite the variability in the behavior, with the path taken by the mice to goal distinct in each trial, the FSA event occurred reliably in most serial and spatial search strategy trials and coincided with the phase of spatial navigation where mice approached the edge of the maze from the central starting point.

Importantly, the sequence of state changes preceding and succeeding the frontal activation event were distinct for goal and non-goal oriented (spatial and serial) search strategies. In spatial (goal-oriented) trials, the 1s period prior to the frontal state activation event (FSA) was marked by a transition from activation of posterior regions of the cortex to broad activation of the lateral regions of the cortex, anterior to the primary visual areas (State 3). In serial (non-goal-oriented) trials, this 1s period was marked by a sequential progression of states associated with high level bilateral activation of posterior regions of the cortex along with the RSC (States 4 and 5), followed by broad activation of left hemisphere (State 7). These distinct sequences of state transitions are summarized in **Figure 3i**. This points to different brain wide circuits being recruited at different time points during the task. Further, these data suggest a frontal role in moving towards the edge and suggests differences in information processing between spatial and serial strategies, both of which successfully get the mouse to the goal.

Goal-oriented spatial navigation depends on cognitive maps ^39^, dependent on structures such as the hippocampus (HPC) and connected cortical circuits ^1,3,40–42^. Recent work looking simultaneously at mesoscale cortical activity and HPC electrophysiology has established a temporal link between mesoscale cortical activity and hippocampal oscillatory such as slow gamma activity and sharp wave ripples in the HPC ^25,43–45^. Such studies confirm previously elucidated systems across hippocampus and cortical brain regions that mediate spatial navigation ^46,47^.

We posit that the distinct spatio-temporal sequences of cortical activation we observed in this study may be part of a larger cortico-hippocampal network computation wherein incoming sensory information seeds retrieval of encoded memory in the HPC followed by reactivation of trace memory in the cortex, followed by execution of motor sequences in which frontal regions of the cortex are active (**Fig. 5**). These sequences are different depending on whether the navigation strategy involves orienting towards a known spatial goal before making an approach or part of a simpler serial search process.

**Figure 5:**
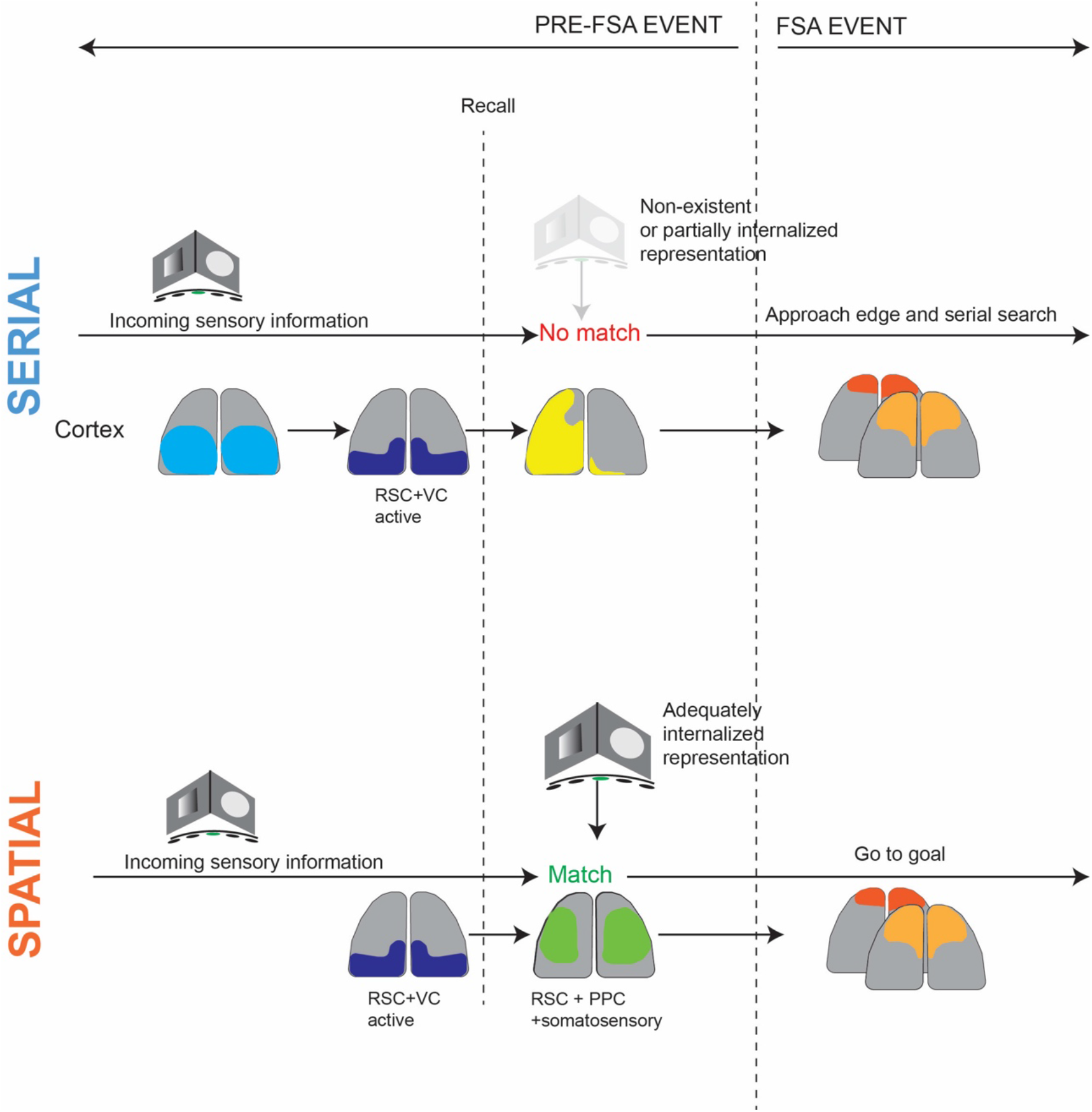
Proposed model for distinct cortical dynamics corresponding to non-goal and goal directed search strategies

## METHODS

### Surgery

Eight Thy-GCaMP6f mice were used in this study ^33^. All animal procedures were performed in accordance with the University of Minnesota’s Institutional Animal Care and Use committee (IACUC). Mice were pre-emptively administered 1 mg/kg slow-release Buprenorphine (Bupenorphine-SR, ZooPharm) and 1 mg/kg Meloxicam prior to surgery. They were then anesthetized using 1-4% isoflurane in pure oxygen prepared for surgery following standard aseptic procedures – the scalp was shaved and sterilized with repeated, alternate scrubbing with Betadine and 70% ethanol. The eyes were covered in sterile eye ointment (Puralube, Dechra Veterinary Products) to prevent drying. The mouse was affixed in a cranial microsurgery robot ^48,49^ under 1-2% isoflurane. The surgical robot performed a large bilateral craniotomy spanning most of the motor, somatosensory, association, higher visual and visual cortices. The top edge of the craniotomy was always cut at 2 mm anterior to Bregma to facilitate alignment of the field of view to the Allen brain atlas ^50^. A transparent polymer skull^51^ compatible with a miniaturized head mounted device ^32^ was initially glued to the skull using surgical grade cyanoacrylate glue (Vetbond, 3M). Two bone screws were implanted in the parietal bone to further anchor the implant, prior to cementing with dental cement (Metabond, Parkell Inc). Mice were recovered from surgery on heating pad and returned to their home cage once they were full ambulatory.

### Barnes Maze behavior

Mice recovered for at least 3 days after surgery. Mice were handled for 15-minute sessions over three successive days prior to experiments to acclimatize them to the experimenter. Dummy mini-mScopes with matched weights to the device used for trials were attached to the mice head to acclimatize them to the device weight during the handling sessions. An experimenter lowered the mice to the center of the maze at the beginning of each trial. Trials were split into three groups, habituation phase, acquisition phase, and probe phase, derived from canonical Barnes maze procedure (Barnes et al. 1979, Pitts et al 2018). During the habituation trail phase, animals were placed in a cylinder in the center of the maze and a dummy mini-mScope was fitted to their implants. Non-goal holes were covered, revealing only the goal hole, and the mouse was allowed to explore the maze for 4 minutes. The maze was then rotated by 90° degrees for acquisition trials. During acquisition trial days, the mini-mscope was fitted onto mice for recording. Mice were placed in the start cylinder in low red-light conditions. Immediately when the trial began, white noise was played at 60 dB and a yellow overhead light was turned on. Non-goal holes were 1 cm deep with black silicone floors. Trials were terminated when the mouse entered the goal hole or after a 3-minute experiment time. For the probe trials, the maze was rotated so that the goal was in a different location with respect to the visual cues. Following all trials, the mouse was placed in the goal box for 1 minute, then returned to their home cage outside of the behavioral enclosure. In between trials, the maze was cleaned with 70% ethanol to reduce odor trails.

The Barnes maze was constructed from a 2.5. cm thick white, high-density polyethylene (HDPE) sheet. A 1-meter diameter circle was cut out of the HDPE sheet. Twenty 10 cm diameter holes were cut into the perimeter 5 cm from the edge of the sheet. A custom-made stair-case goal box was 3D printed using 1.75 mm diameter black PLA filament on a fused deposition modeling 3D printer (M2 3D printer, MakerGear). The maze was mounted onto an aluminum extrusion frame and anchored to a behavioral enclosure. The maze was 0.6 meters from the ground and at least 1.5 meters from any wall. The walls of the behavioral enclosure were made from 1/8-inch-thick single plywood sheets (Eucatile white tile board, Home Depot) and were coated with acoustic damping foam on the inner walls (JBER Acoustic Sound Foam Panels, Amazon) that covered the 1.8 m x 1.8 m x 2.4 m enclosure. A single behavior camera was mounted 1.2 m above the center of the arena to record behavior during the experiments (Blackfly S USB-3, FLIR). The mini-mScope electronics were routed through a low torque commutator (Carousel Commutator 1x DHST 2x LED, Plexon Inc).

### Cortex-wide imaging using mini-mscope

#### Behavior imaging

One overhead camera was used to capture the entirety of the Barnes maze. The behavior camera was set to external trigger mode, line 3 trigger, any edge, (Spinview) and was synchronized to capture frames with the TTL pulses sent by the mini-mScope at each frame capture. The behavior camera exposure was set to 1000 μs and the resulting frames were compressed by 25% and saved to random access memory (128 GB RAM) as a .avi video file.

#### Calcium imaging

The original mini-mScope CMOS sensor ^32^ was replaced with the MiniFAST CMOS sensor (Sony IMX290LLR-C CMOS sensor, Framos) for its increased sensitivity and smaller pixel size (Juneau et al. 2020). The MiniFAST sensor was set to acquire images at 30 frames per second (FPS), with each frame alternating between blue and green light illumination. Thus, images were acquired at 15 FPS under each illumination condition. The CMOS gain was set to a value of 55, and the LED voltage and current for the green LEDs was 5V 0.2A and 8V 0.8A for the blue LEDs. The blue and green LEDs on the mini-mScope were pulsed for 120 seconds, prior to the experiment to allow them to warm up and reach a stable intensity. The mice were brought into the Barnes maze under red light and placed into the opaque cylinder at the center of the maze ~90 s after the LEDs were turned on. The mini-mScope was attached to the mice via 3 interlocking magnets. At ~120 seconds, the white noise and yellow LEDs in the Barnes maze were switched on and the opaque cylinder was removed, marking the start of the trial. Trials typically lasted until mice went into the goal hole or at the end of 180 seconds.

#### Data pre-processing

##### Behavior data pre-processing

For each trial, the location of each hole in the Barnes maze and the outer shape of the maze was automatically detected using computer vision scripts to define regions of interests (ROIs) within the Barnes maze. The location of the goal hole was marked to track where it was located, as the Barnes maze was rotated across cohorts and probe days. The behavior camera data was aligned with the calcium imaging data via timestamps generated by the CMOS data acquisition board. Any frame drops or motion artifacts detected in the calcium imaging data were dropped in both the calcium imaging data and the behavior imaging data. The behavior camera data was also down sampled to match the calcium channel from the mini-mScope.

##### Calcium data pre-processing

To assist with data saving, the MiniFAST software saves calcium imaging data in separate 1000 frame videos. The individual 1000 frame videos were combined into a single video using custom MATLAB scripts (2022b, MathWorks). The mean pixel intensity of each frame was calculated, and K-means clustering was used to classify each mean pixel intensity of the video into the blue and green channels. Frames that were not classifiable into either the blue nor green channels due to large motion artifacts or irregularities in LED intensity (~0.04% of all frames) were marked for removal in future analysis. The videos corresponding to both illumination wavelengths were then passed through a motion correction algorithm ^53^.

The calcium data videos were compressed to 80% of their original size with a bilinear binning algorithm (2022b, MathWorks). One frame randomly selected in each trial was used to draw a mask around the imaged brain surface and exclude the background and superior sagittal sinus artery to reduce noise in the overall DF/F signal. For each mouse, the masks across all trials were averaged to generate a mouse-specific average cortex mask. The average mask was imposed across images acquired in all trials for a mouse so that the number of pixels used in each analysis remained consistent.

Each pixel within the mask was corrected for global illumination fluctuations using a correction algorithm that produces DF/F data^54^. The DF/F data was filtered using a zero-order phase Chebyshev band-pass filter with cutoff frequencies of 0.1 Hz and 5 Hz (2022b, MathWorks). The resulting data was then spatially filtered with a 7-pixel nearest-neighbor average using a custom MATLAB (2022b, MathWorks) script. The resulting DF/F time series for each pixel was then z-scored.

### Data Analysis

#### Behavior

Data from the overhead behavior camera was analyzed using an unsupervised, marker-less tracking algorithm (DeepLabCut^55^). The program was trained to track the nose, the top of the head/mini-mScope, between the ears, the right and left forepaws, the shoulder blades, right and left hind paws, the lower back, the base of the tail, and the tip of the tail. This tracking data was used to determine where the mice were in the Barnes maze throughout the trial. To classify search strategy, the Barnes maze was split into 4 equal quadrants and each hole was automatically detected and labeled. Random trials were classified if the mouse’s tracking trajectory crossed over 3 quadrants of the maze non-sequentially before reaching the goal. Serial trials were classified if the mice traveled less than 3 sequential quadrants and covered at least 3 sequential holes on either end of the goal hole. Spatial trials were defined if the mice traversed less than 2 sequential quadrants and no more than 1 sequential hole on either side of the goal hole. Radially, the maze was divided into the central circle, the approach zone and the serial exploration zone, with the diameter of the central circle corresponding to the length of the mice (60 pixels), and the inner radius of the serial exploration zone being one length of the mouse lesser than the outer diameter of the maze.

#### State identification using image correlation clustering

All calcium data was analyzed using custom scripts in MATLAB (2022b, MathWorks). At each time point, the DF/F z-score for the current frame was correlated with all frames across trials per mouse using a Pearson’s correlation coefficient to construct a correlation matrix across trials (**Figure 2b**). The correlation matrix was then sorted using k-means clustering with RNG defaults for reproducibility and with 5000 maximum iterations and 500 replicates to search for common, reoccurring activity patterns across time. A t-distance optimization algorithm was used to determine the optimal number of clusters to sort the correlation matrix, so that the correlations within each cluster were maximized and correlations across clusters were minimized ^34^. The number of clusters for which the largest cumulative t-distance value obtained was selected as the number of clusters or states for each mouse (**Supplementary Figures 1-2**). All the frames within an identified cluster were averaged to generate a mean activity spatial map for each state. Image correlations between these mean activity maps for each state identified for all mice were computed to construct a second correlation matrix, which was then sorted into 7 clusters via k-means clustering (**Figure 2c, Supplementary Figure 1a, Supplementary Figure 2**).

#### Frontal state activation

The time series of state activations for all trials were filtered using a sliding window to extract periods of high activation of state 1 and 2 (the frontal states) for all trials. The frontal state activation event was determined to be present if it persisted for a period greater than 1 second, with up to 4 frames of jitter into other states before returning to state 1 or state 2. After the events were labeled, all state activation time series were aligned to the start and end of the frontal state activation event period for statistics and further analysis.

#### Head orientation angle

Two angles were defined for head orientation of the mouse during the start of the trial until the frontal state activation period. The allocentric angle, denoted as ω, was the angle between the instantaneous mouse body-head vector relative to fixed vector drawn from the center of the maze to the goal. The egocentric angle, denoted as φ, was the angle difference between the instantaneous mouse body-head vector and vector drawn from the instantaneous position of the mouse’s body to the goal location.

#### Statistics

Wilcoxon rank sum non-parametric tests were used to determine the statistical significance between serial and spatial search strategies’ state activation (**Figure 3e, Figure 4 h,l**). A Kruskal-Wallis test was used to determine statistical significance between head direction angles in the pre-FSA period. Non-parametric tests allow for unequal sample sizes between the search strategies. ANOVA tests were run with a Bonferroni correction to determine the significance of state activation in the peri-event state probability histograms (**Figure 2g**). All error bars denote standard deviation.

## ACKNOWLEDGEMENTS

SBK acknowledges support from National Institutes of Health grants R01NS111028, RF1NS113287, 1RF1NS126044, P30DA048742 and the McKnight Foundation. MLR was supported the University of Minnesota Informatics Institute Graduate Fellowship in 2021, and the University of Minnesota Institute of Engineering in Medicine (IEM) Graduate Fellowship in 2022. ADR acknowledges support from R01MH112688. We thank Dr. Tim Buschman and Ryan Peters for helpful comments on the manuscript.

## AUTHOR CONTRIBUTIONS

DS and MLR contributed equally. DS, MLR, SBK designed experiments, DS and MLR conducted the experiments. DS, MLR, EK, KS, ADR and SBK performed data analyses, DS, MLR, EK, KS, ADR and SBK wrote the manuscript.

