## Supplementary Figures for "Distinct mesoscale cortical dynamics encode search strategies during spatial navigation"

**SUPPLEMENTARY MATERIAL**

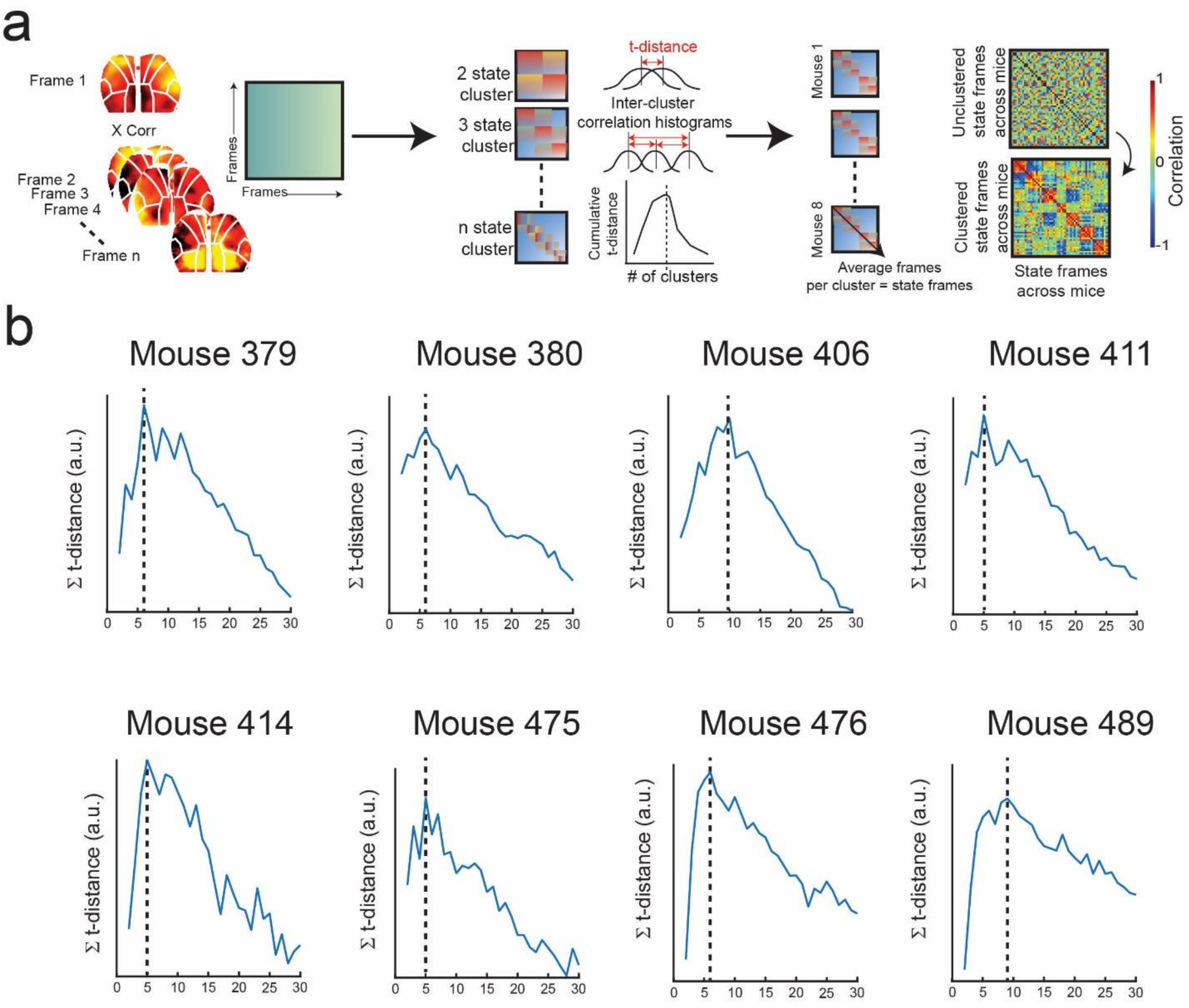


**Supplementary Figure 1: Methodology for identifying brain states a)** *Left:* An image correlation matrix is constructed for all frames across all trials for each mouse. *Center:* A t-distance optimization algorithm is used to determine the optimal number of states to cluster the image correlation matrix into via k-means clustering for each mouse. Since the number of clusters (defined as states here) is unknown apriori, the t-distance optimization maximizes the correlation within each state while minimizing the correlation across states. The maximum t-distance value indicates the optimum state number for that mouse. *Right:* Mean DF/F activity maps for each state are constructed by averaging all the frames identified in that state for each mouse. A second image correlation matrix is formed by taking the correlation between the mean DF/F activity maps that exist for each optimized state in each mouse. The correlation matrix is sorted into 7 states of highly correlated average DF/F activity maps across mice, forming a common state model across mice. **b)** T-distance plots for the optimum number of states found for each mouse using the algorithm described in **a.** There are 5-10 optimum states found across n=8 mice.

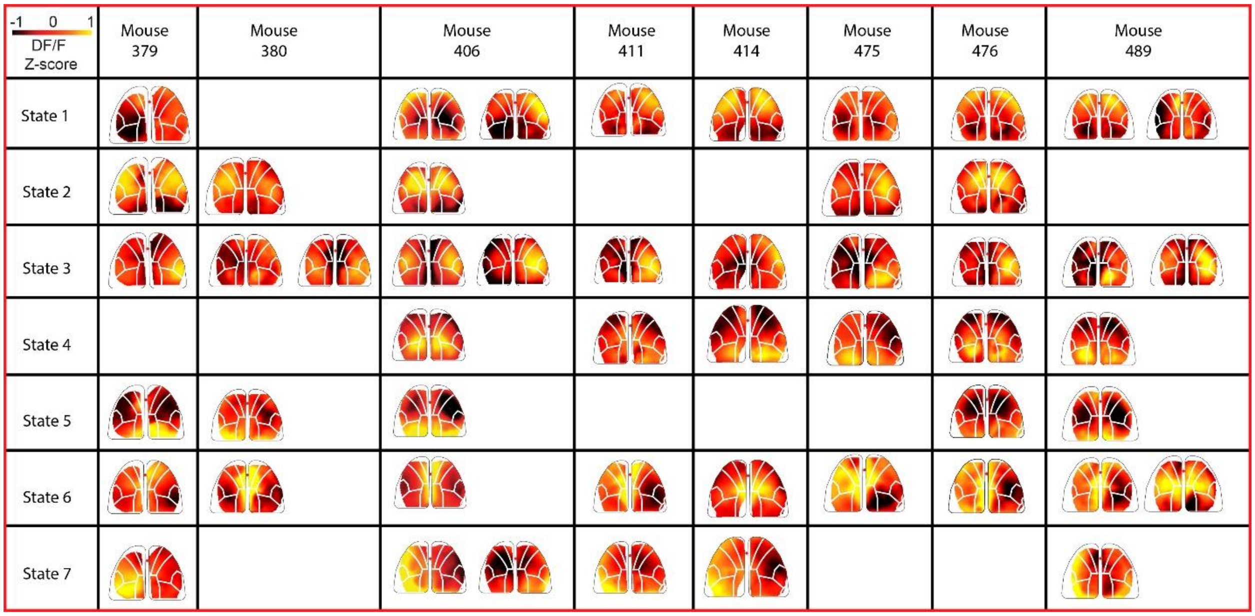


**Supplementary Figure 2: Common state space across mice.** Mean DF/F z-score activity maps are formed for each mouse and state by averaging the DF/F activity across all frames identified within each state. The average maps are sorted into the common 7-state model and tabulated here (see **Supplementary Fig. 1a** for details).

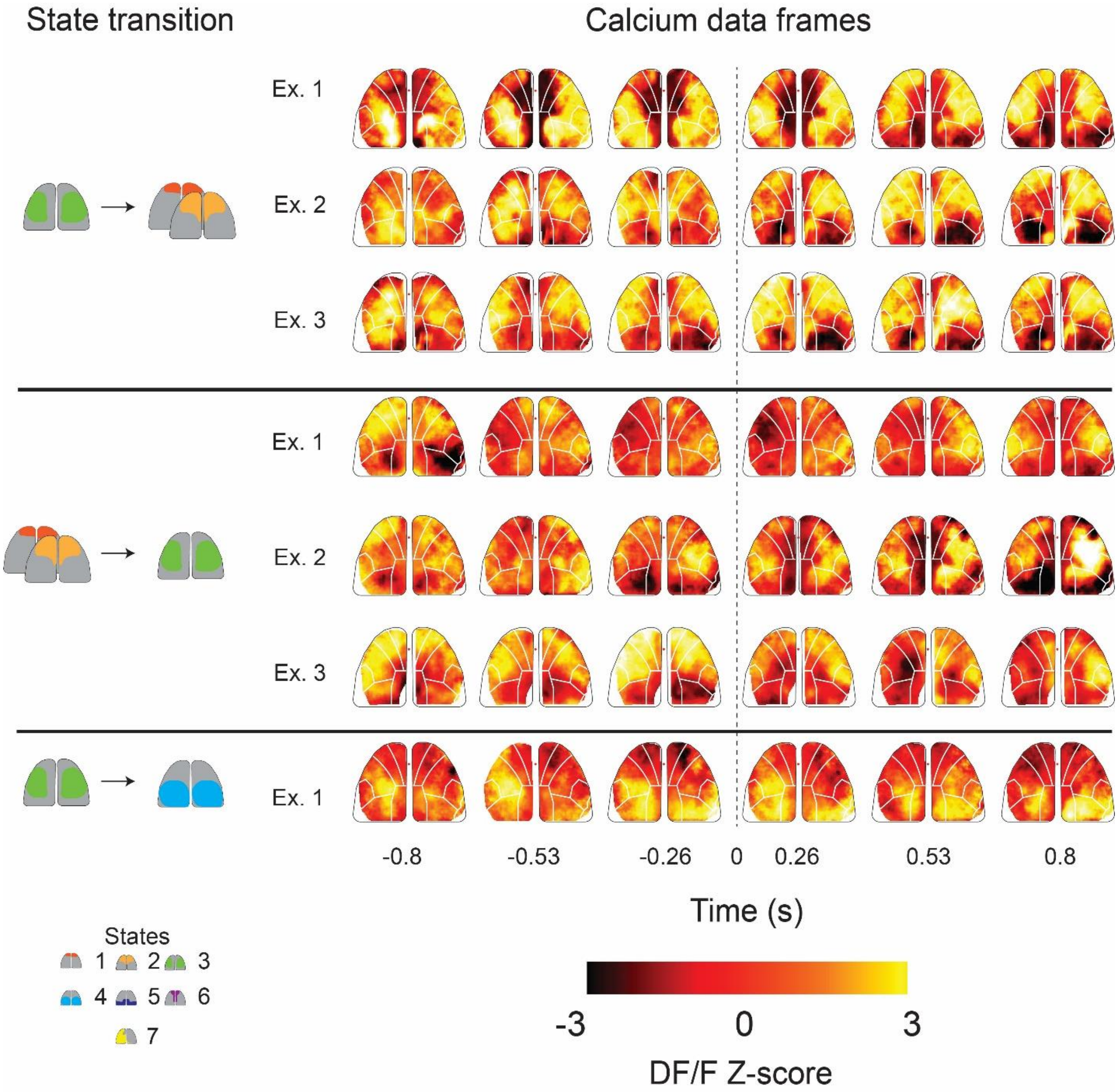


**Supplementary Figure 3: Montages of changes in calcium activity across the cortex during common state transitions.** DF/F z-score calcium activity maps illustrating examples of common state transitions to and from state 3. Simplified representations of the transitions are on the left, and DF/F z-score maps are on the right, with each row representing a different example. Example transitions occur at t = 0. *Top:* three examples showing transitions from frontal states 1 and 2 to state 3. *Middle:* three examples showing transitions from state 3 to frontal states 1 and 2. *Bottom:* one example of a state 3 to state 4 transition.

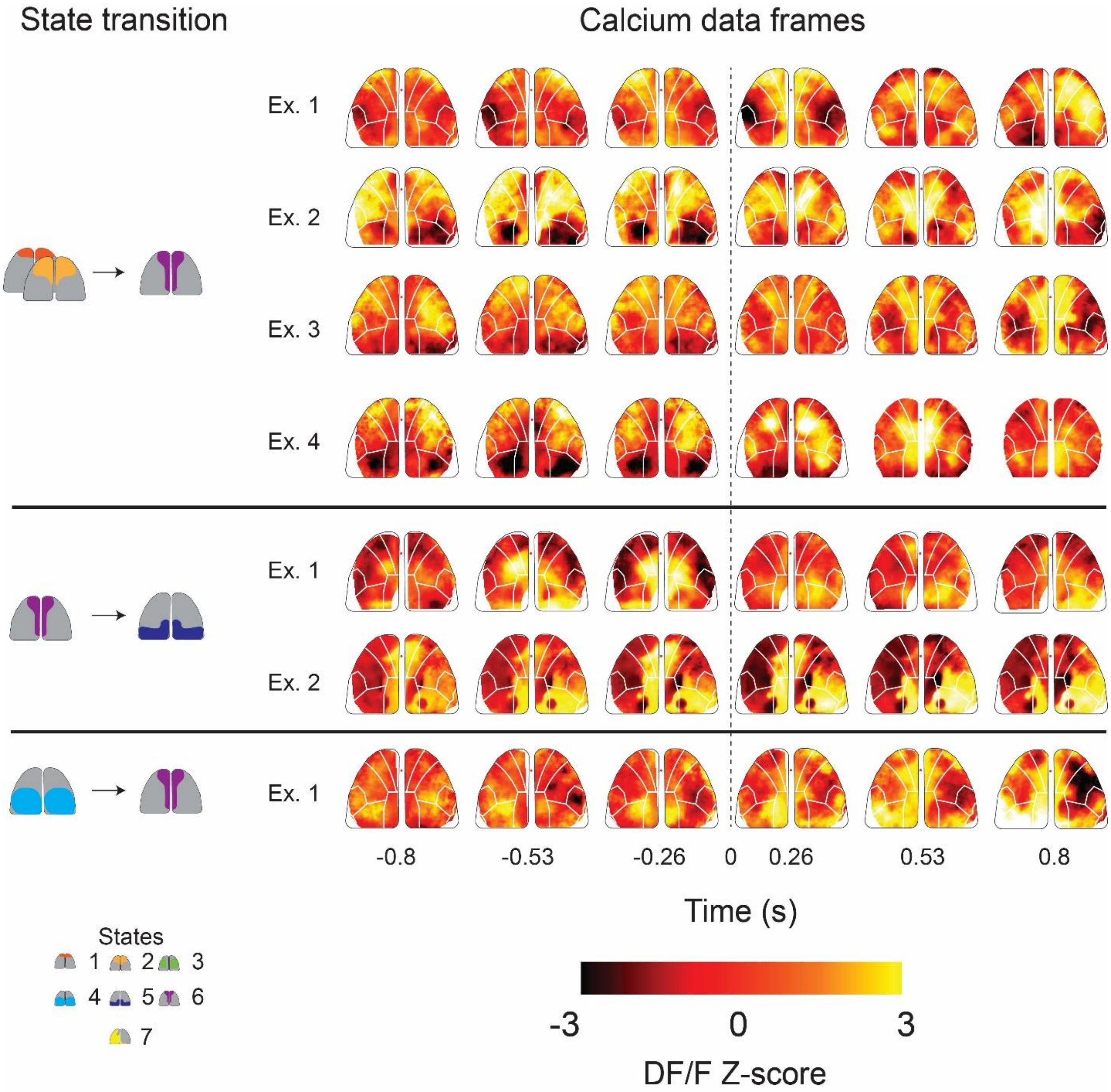


**Supplementary Figure 4: Montages of changes in calcium activity across the cortex during common state transitions.** DF/F z-score calcium activity maps illustrating examples of common state transitions to and from state 6. Simplified representations of the transitions are on the left, and DF/F z-score maps are on the right, with each row representing a different example. Example transitions occur at t = 0. *Top:* four examples showing transitions from frontal states 1 and 2 to state 6. *Middle:* two examples showing state 6 transitioning to state 5. *Bottom:* one example of a state 4 to state 6 transition.

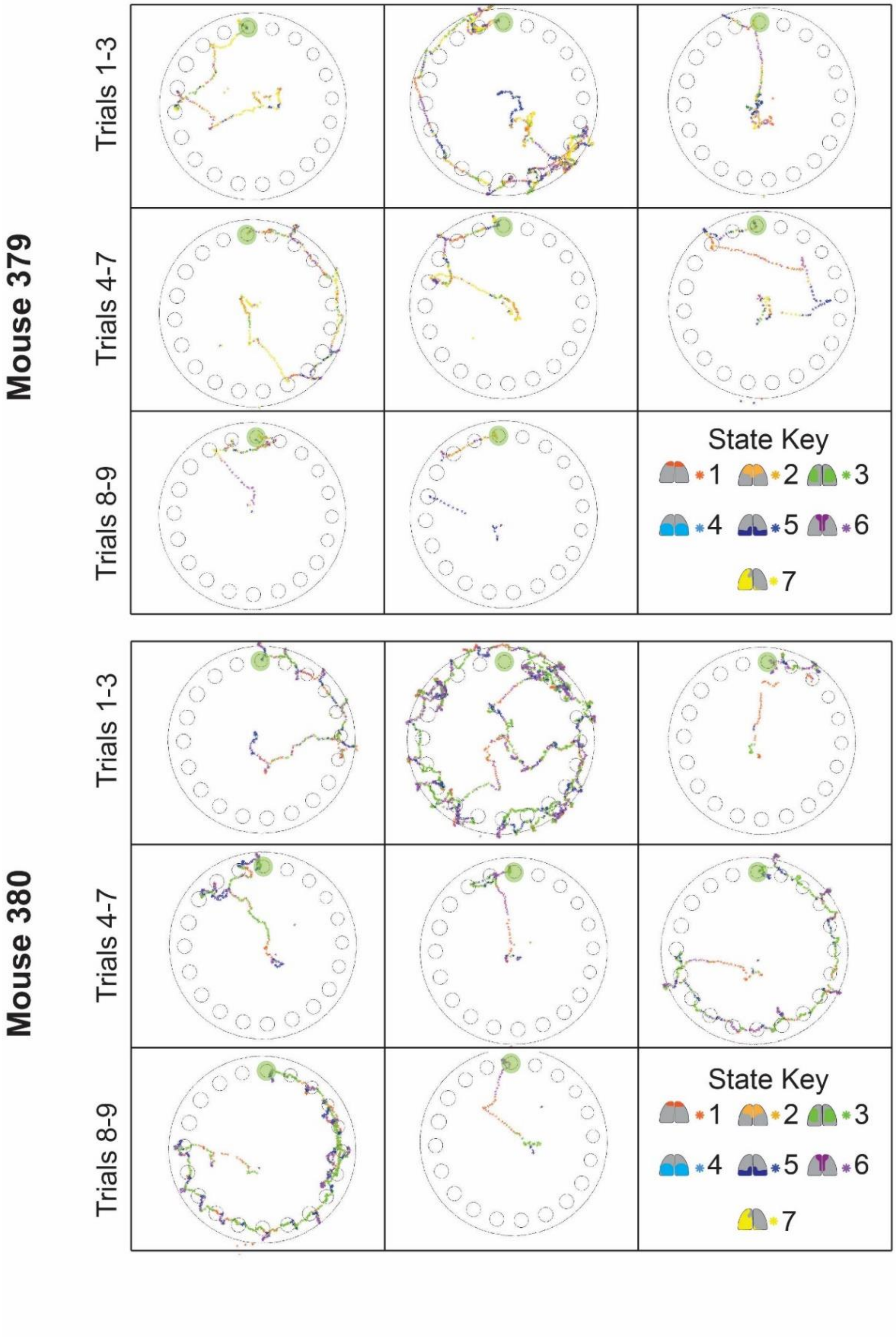


**Supplementary Figure 5: State activation along mice paths.** State activation plotted along paths taken by mice for all acquisition trials are shown for mice 379 and 380**.**

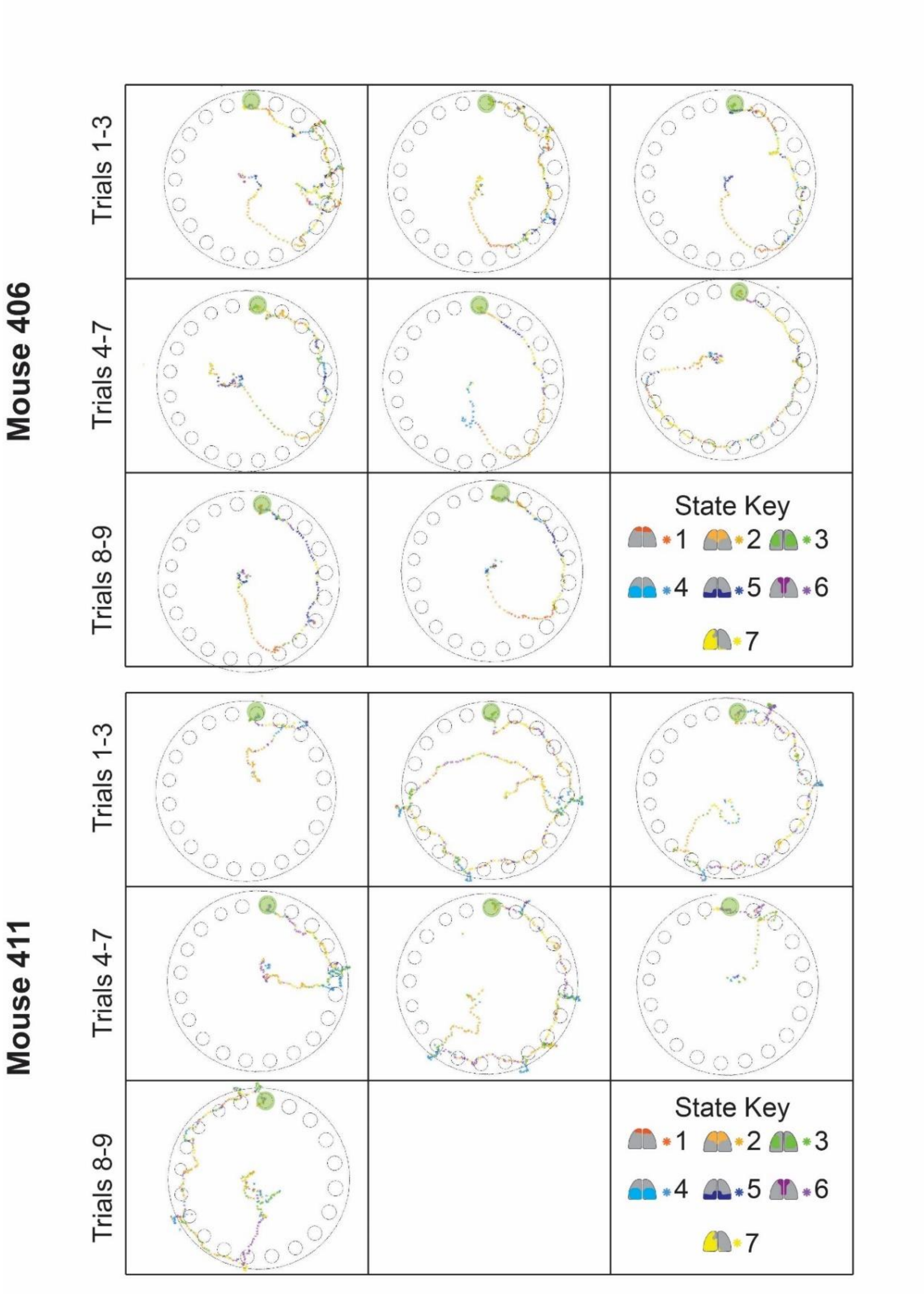


**Supplementary Figure 6: State activation along mice paths.** State activation plotted along paths taken by mice for all acquisition trials are shown for mice 406 and 411.

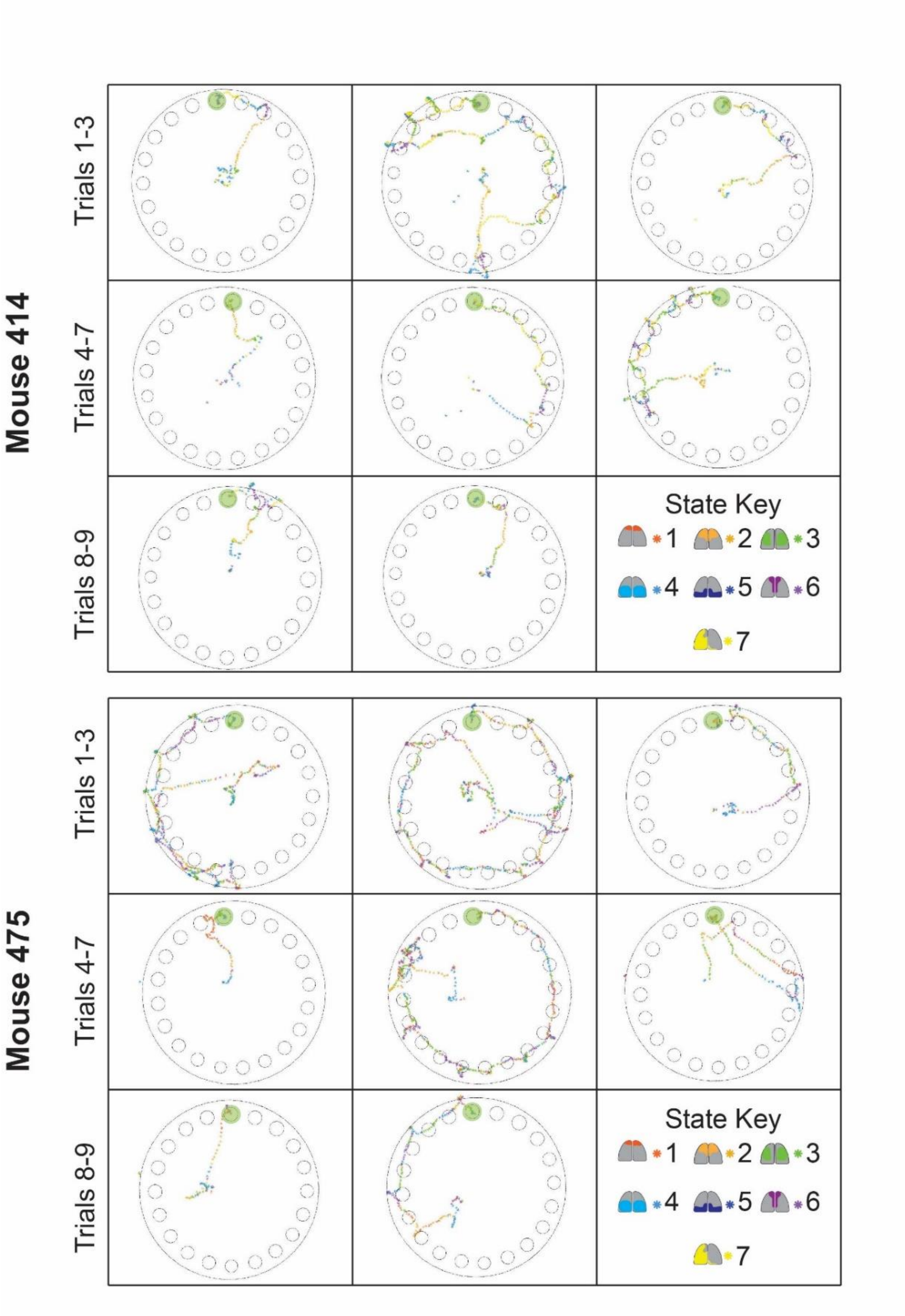


**Supplementary Figure 7: State activation along mice paths.** State activation plotted along paths taken by mice for all acquisition trials are shown for mice 414 and 475.

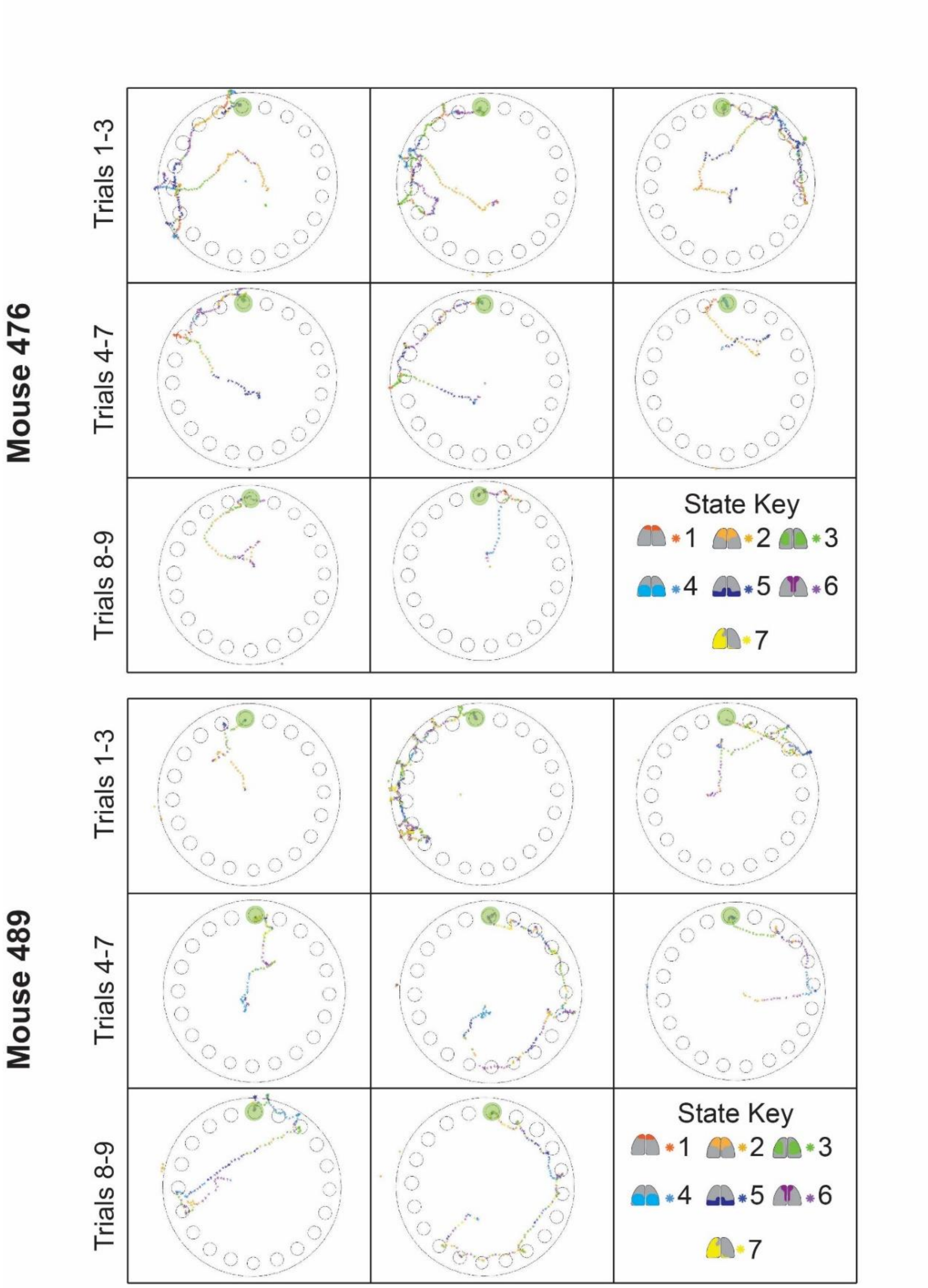


**Supplementary Figure 8: State activation along mice paths.** State activation plotted along paths taken by mice for all acquisition trials are shown for mice 476 and 489.

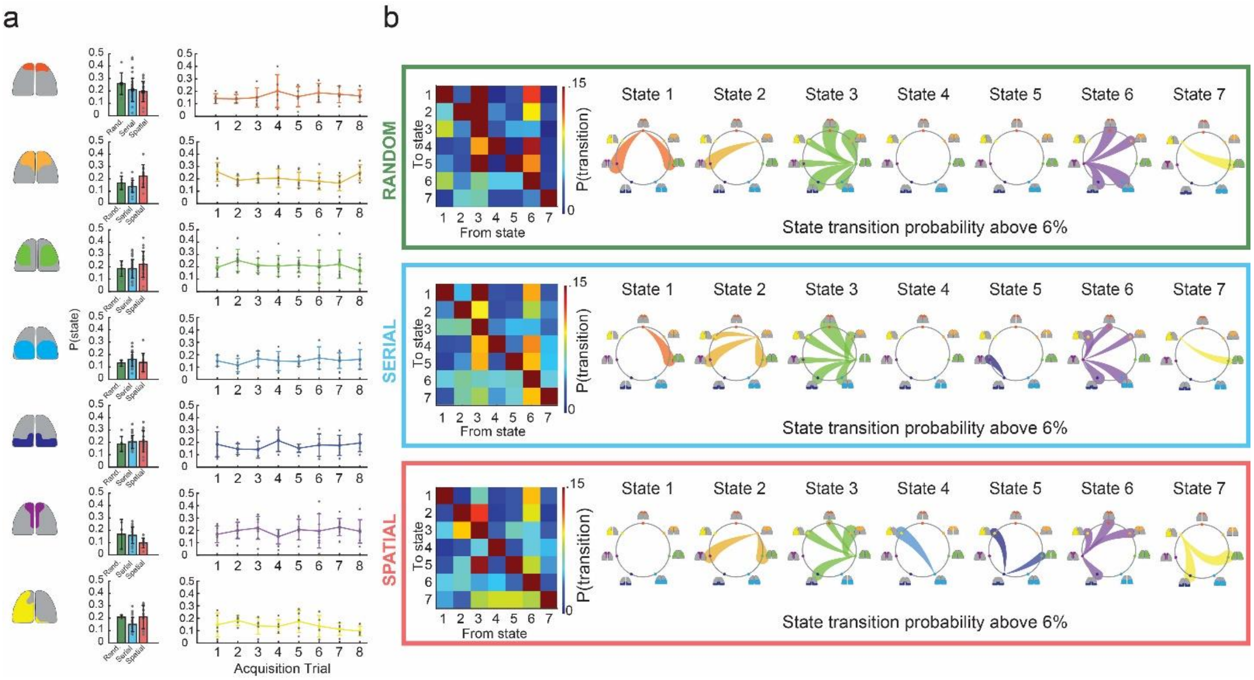


**Supplementary Figure 9: Brain-state variations across trials and search strategies**. **a)** State activation probabilities across trials and search methods. *Left:* simplified schematic for each state. *Center*: state activation probabilities for trials groups by search method. Data points indicate each trial which mice utilized the search method. *Right:* state activation probabilities across trials for all mice. Data points indicate state activation probabilities for each mouse. All error bars indicate standard deviation. **b)** State transition probability matrices for random (top), serial (middle), and spatial (bottom) trials. State transition node maps indicate specific state transition probabilities above 6% for each search method. Thickness of the lines indicate higher state transition probabilities.
